# Task relevance selectively modulates sensorimotor adaptation in the presence of multiple prediction errors

**DOI:** 10.1101/2024.10.07.616966

**Authors:** Somesh N Shingane, Nishant Rao, Neeraj Kumar, Pratik K Mutha

## Abstract

Adaptation to consistently occurring sensorimotor errors is considered obligatory in nature. We probed the robustness of this finding by asking if humans can selectively attenuate adaptation based on the task-relevance of error signals. Subjects made planar reaches to three different targets: an arc (Experiment 1), a bar (Experiment 2), and a point (Experiment 3). During the reach, perturbations in extent (visuomotor gain), direction (visuomotor rotation) or both simultaneously were employed. In Experiment 1, subjects showed robust adaptation to the rotation when reaching to the arc even though the presence of this perturbation was irrelevant for achievement of the task goal. Interestingly however, rotation adaptation was strongly attenuated when it was presented simultaneously with a task-relevant gain perturbation. In Experiment 2, which involved reaches to the bar, again, subjects successfully adapted to the task-irrelevant gain perturbation when it occurred in isolation. However, adaptation was attenuated when the gain co-occurred with a task-relevant rotation. Experiment 3 revealed that the attenuation observed in the first two experiments was not due to an inability to adapt to co-occurring rotation and gain perturbations. Collectively, our results suggest that the sensorimotor system selectively tunes learning in the presence of multiple error signals, a finding that can potentially be explained by a biased competition mechanism. That is, given limited processing capacity, a salient attribute – the relevance of the error to the task goal in this case – is prioritized for processing and drives subsequent adaptive changes in motor output.

**NEW AND NOTEWORTHY:** The motor system continuously uses error feedback to recalibrate movements in response to changes in body and environmental conditions. Such error-based adaptation is thought to be obligatory, occurring whenever error signals are present, and even if the learning interferes with achievement of the task goal. Contrary to this classical view, we demonstrate selective modulation of motor adaptation in the presence of multiple error signals based on their task-relevance.

## INTRODUCTION

Motor adaptation is the process through which humans adjust their movements in response to errors induced via destabilizing environments. Adaptation ensures that movements remain well-calibrated and accurate despite changing task conditions. Adaptation to environmental and also bodily changes has been demonstrated for a wide variety of actions including eye movements (Barne et al., 2023; Tian et al., 2009; Tyralla et al., 2023), point-to-point reaching and grasp-and-lift movements (Hadjiosif et al., 2023; Krakauer & Shadmehr, 2006; Parikh et al., 2020; Rao et al., 2021; Shadmehr et al., 2010) and locomotion (Park et al., 2022; Torres-Oviedo et al., 2011). It is increasingly appreciated that motor adaptation is driven to a great extent by sensory prediction errors, or the difference between the expected and actual sensory consequences of movements. Such learning is thought to bring about implicit refinements to motor plans, and is distinct from the effects of deliberative strategies that may be employed to reduce errors induced by the perturbing environment (Cressman & Henriques, 2015; Hegele & Heuer, 2010; Inoue et al., 2015; Morehead et al., 2017; Oza et al., 2024; Sadaphal et al., 2022; Taylor et al., 2014; Tsay et al., 2022).

Some past work suggests that the sensorimotor system does not respond equally to all prediction errors (Kasuga et al., 2013; Wei & Körding, 2009). Sensitivity to error, measured as a change in motor output on a movement following experience of a perturbation on a previous movement, appears to be higher for smaller prediction errors (attributed to the body) than larger ones (attributed to the environment). Yet, despite this robustly demonstrated tuning of the adaptive response, it is generally believed that adaptation cannot be entirely suppressed. In fact, it has been shown that prediction error driven adaptation occurs even when it is detrimental to success in the task (Mazzoni & Krakauer, 2006). Other investigations have shown that adaptation cannot be attenuated by instruction (Bond & Taylor, 2015; Kim et al., 2019; Morehead et al., 2017; Oza et al., 2024; Tsay et al., 2022) and that it occurs even when prediction errors are completely irrelevant to the task and there is no need to compensate at all (Morehead et al., 2017; Schaefer et al., 2012).

In an elegant study, Schaefer et al. (2012) had subjects reach to an arc-shaped target and used a visuomotor rotation to introduce errors between the expected and actual hand position displayed as a cursor on a screen. Because the target was an arc, the rotation in no way affected achievement of the task goal of bringing the cursor to the target; subjects therefore did not need to compensate for the rotation on a trial-to-trial basis. Yet, subjects showed clear evidence of adaptation. Similar inferences could be drawn from the work of Synofzik et al. (2006), who showed robust adaptation to a visuomotor rotation introduced when movements were made even without a precise target location being defined. These findings reinforce the idea that the motor system cannot tolerate the occurrence of prediction errors and always attempts to cancel them.

The goal of the current study was to test the limits of this obligatory nature of adaptation. Specifically, we asked whether participants would continue to adapt to a task irrelevant perturbation even when it co-occurred with a task relevant one. We addressed this by simultaneously imposing rotation and gain perturbations that introduce errors in direction and extent respectively and manipulating their relevance for achievement of the task goal. It must be pointed out that our aim was not to establish whether the motor system would adapt to two simultaneously imposed perturbations; Kasuga et al. (2013) earlier and Tsay et al. (2024) very recently, have nicely demonstrated that adaptation in the presence of two simultaneously occurring but similar error signals (cursor rotations) is indeed possible. Rather, our attempt was to challenge the notion that adaptation cannot be selectively modulated in the presence of prediction errors. We show that motor adaptation can in fact be constrained by manipulating the relevance of prediction errors, which suggests that such learning is not always obligatory and can be flexibly tuned.

## METHODS

### Subjects

105 young, neurotypical, right-handed individuals (age range: 18-40 years) participated in this study that comprised of 3 different experiments. Subjects did not report any neurological impairment, musculoskeletal disorder or physical injuries and provided written informed consent to participate in the experiment. The study was approved by the Institute Ethics Committee of the Indian Institute of Technology Gandhinagar and study procedures followed the principles expressed in the Declaration of Helsinki.

### Apparatus and general task procedures

Subjects sat facing a digitizing tablet (Calcomp Inc) integrated with a virtual reality setup. The setup consisted of a horizontally mounted HDTV that projected onto a mirror mounted horizontally between the TV and the tablet (Figure 1A). Subjects could not see their arms since the mirror blocked their view; they performed the task while looking into the mirror, which reflected the task elements displayed on the TV screen above.

**Figure 1:**
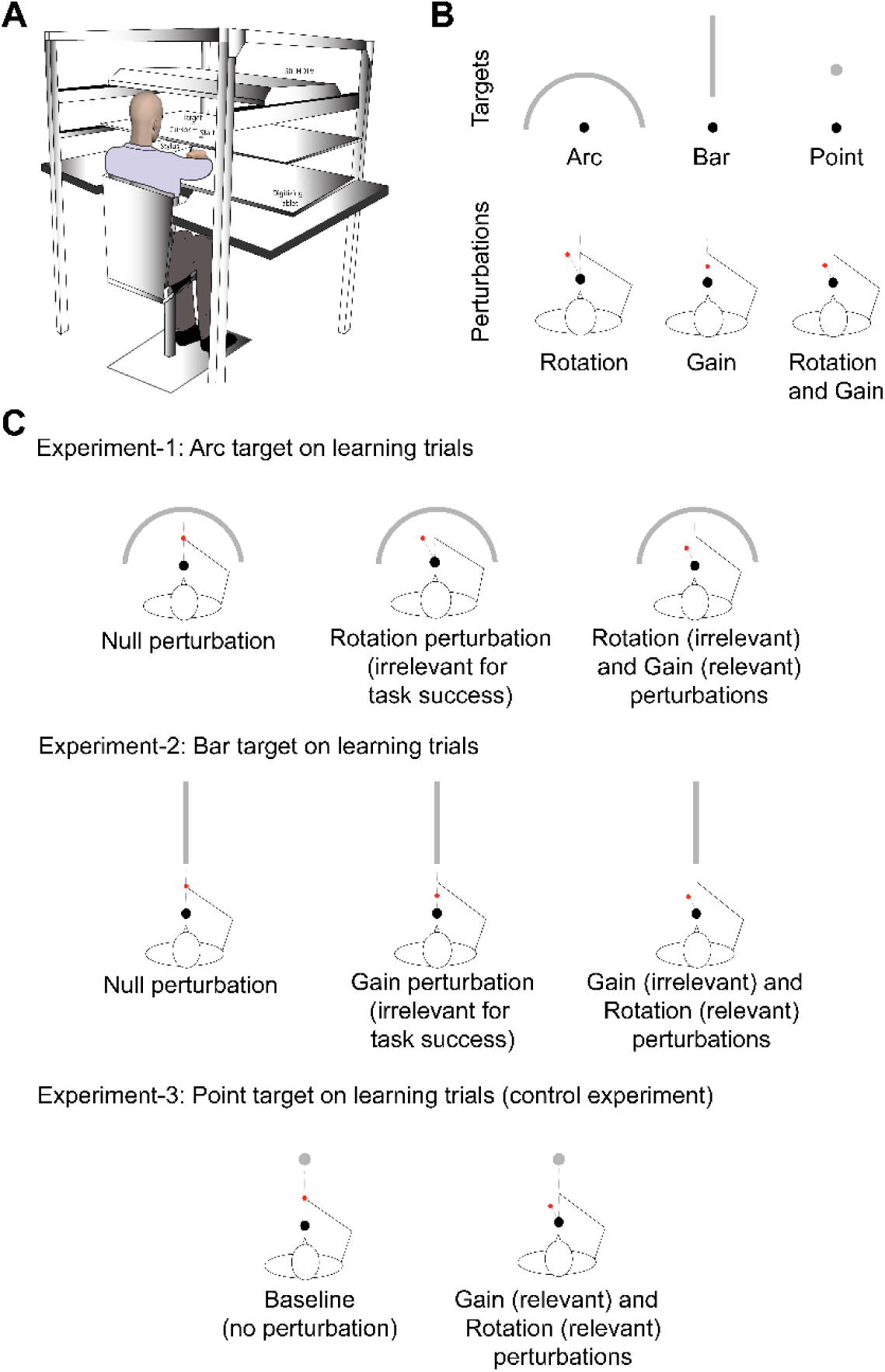
Experimental design and setup. (A) Subjects sit at a table and make planar reaches on a digitizing tablet mounted on it using a stylus held in their right hand. Visual feedback about the motion is projected as a cursor on a screen suspended above the tablet. A mirror that reflects the on-screen display is placed between the tablet and the screen. (B) Targets and perturbations used in the study. Broadly, three different targets were used: an arc, a bar and a point. These could be paired with different perturbation types, a visuomotor rotation, a visuomotor gain and a rotation and gain imposed simultaneously (rotation+gain). (C) Target-perturbation combinations. When reaching to the arc target, different groups of subjects were exposed to either a “null”, a rotation or a rotation+gain perturbation. Likewise, when reaching to the bar target, subjects experienced null, gain or rotation+gain perturbations. Finally, for the point target, subjects experienced the rotation+gain perturbation only (rotation perturbation in point rotation+gain condition was actually applied clockwise but shown in the figure as counterclockwise for homogenous visualization with previous other experimental conditions).

Subjects were instructed to make quick planar reaching movements from a single circular start location (0.62 cm radius, white color) to a target while holding a stylus in their right hand (Figure 1B, top panel). The target could be either a semi-circular arc (12 cm radius, 0.62 cm thickness, center coinciding with the start position, white color), a bar (12 cm length, 0.62 cm thickness, beginning 6 cm from the start position, white color) or a point (0.62 cm radius, 12 cm away from the start position, white color). For the arc, subjects could move in any direction and still be successful in the task as long as they moved the right extent. The direction dimension was thus irrelevant during arc reaches and only the extent dimension mattered. In contrast, for the bar, subjects could move any extent but had to reach specifically along the forward direction to hit it. Thus, movement extent was irrelevant for the bar target, but movement direction was highly relevant. For the point target, naturally, both direction and extent were relevant. During the reach, hand position was displayed by means of a feedback cursor (0.4 cm radius, red color) throughout the movement. Cursor motion could be veridical with the hand or perturbed in various ways depending upon the experimental conditions.

To initiate a trial, subjects brought the cursor into the start position and held it there for at least 500 ms. After this time, the target appeared, and subjects were required to reach to it. Subjects experienced 3 types of trials: baseline, learning and catch. On baseline trials motion of the hand (on the tablet) and the cursor (on the screen) remained mapped to each other. On learning trials, this relationship between hand and cursor motion was perturbed. We used three different kinds of perturbations (Figure 1B, bottom panel): a rotation, a gain, and a rotation and gain employed simultaneously (rotation+gain). Furthermore, on learning trials, we used only a single target (either arc, or bar, or point depending on the experiment; see below). Finally, catch trials were interspersed among learning trials to assay whether subjects were adapting to the perturbations. On the catch trials, subjects always reached to a point target under veridical visual feedback, i.e. no perturbation was applied on the catch trials.

The rotation perturbation was implemented by rotating the motion of the cursor relative to the hand. The gain perturbation was implemented by reducing the extent of cursor motion relative to that of the hand. That is, if the hand moved a certain distance on the tablet, the cursor would move less on the screen. For the rotation+gain perturbation, both these perturbations – the rotation and the minifying gain – were employed together on the same trial. By combining these different perturbations with distinct target types, we were able to manipulate the relevance of the perturbation to achievement of the task goal. Relevance was defined in terms of whether the presence of the perturbation impeded task success or not. For instance, the rotation perturbation would be irrelevant for movements to the arc because subjects would always hit the arc target regardless of which direction they moved (as long as they moved the right extent). In other words, the rotation would not impede goal achievement. However, perturbations in movement extent (gain) would be highly relevant during arc reaches. Likewise, an extent perturbation would be irrelevant for reaches to the bar, but any directional perturbation (rotation) would be highly relevant since, if left uncompensated, it would hinder achievement of the task goal. For the point target, both the rotation and the gain would naturally be task relevant. These combinations would enable us to test how subjects accounted for perturbation relevance, and more importantly, whether they would differentially adapt to the relevant and irrelevant perturbations when they were simultaneously induced.

### Experiment 1: Irrelevant Rotation Perturbation

In Experiment 1, subjects (n = 45) were randomly divided into three groups (n = 15 per group). Subjects in all groups first performed 20 baseline trials in which they reached to either the arc (12 cm radius, 0.62 cm thickness, center coinciding with the start position, 11 trials) or the point (0.62 cm radius, located 12 cm from the start position, 9 trials) target under veridical cursor (0.4 cm radius) feedback. This was followed by 150 learning trials during which they reached only to the arc target under perturbed visual feedback. The first group (ArcNull) experienced a “null” perturbation (essentially no perturbation, veridical cursor feedback), the second group experienced the 30-degree counterclockwise rotation (ArcRot), while the third experienced the simultaneous “rotation+gain” (30-degree counterclockwise rotation, 0.7 minifying gain, ArcRot+Gain). All subjects, regardless of the group to which they belonged, also experienced 10 intermittently embedded catch trials, which, as stated earlier, comprised of reaches to the point target without any perturbation and under continuous cursor feedback. Note that the rotation perturbation was irrelevant as far as the goal of hitting the arc was concerned, but a change in gain (in the rotation+gain combination) would be relevant.

### Experiment 2: Irrelevant Gain Perturbation

The task conditions in Experiment 2 were largely similar to Experiment 1. Briefly, subjects (n = 45) were again divided into 3 groups (n = 15 each). During baseline, all subjects made 20 reaches to either a bar (0.62 cm thickness, length 12 cm, beginning 6 cm from the start position, 11 trials) or a point (0.62 cm radius, located 12 cm from the start position, 9 trials) under veridical cursor (0.4 cm radius) feedback. This was followed by 150 learning trials to the bar target during which the first group experienced the “null” perturbation (BarNull), the second group received the 0.7 times minifying gain (BarGain), and the third group was exposed to the “rotation+gain” (30-degree counterclockwise rotation, 0.7 minifying gain, BarRot+Gain). As in our first experiment, 10 catch trials were embedded within the learning trials. Note that in this case, any change in “gain” was irrelevant to achieving the task goal. In contrast, any perturbation-induced change in direction would be highly relevant and necessitate changes in the actual hand direction. Our interest remained in examining whether adaptation to the irrelevant dimension would be altered in the presence of another relevant perturbation.

### Experiment 3: Control Experiment

In Experiments 1 and 2, one of the dimensions in the “rotation+gain” perturbation (either direction or extent) was always irrelevant for achievement of the task goal. We therefore designed a third, control experiment in which both dimensions would be relevant. In this experiment, only one group of subjects was recruited (n=15). Here, subjects made 45 baseline reaches to all three targets: the arc (12 cm radius, 0.62 cm thickness, center coinciding with the start position, 15 trials), the bar (18 cm length, 0.62 cm thickness, beginning 6.5 cm from the start position, 15 trials) and the point (0.62 cm radius, located 12 cm away from the start position, 15 trials). This was followed by 80 learning trials, during which subjects reached to the same point target under the “rotation+gain” perturbation (30-degree clockwise rotation and 0.6 minifying gain). As before, embedded within these learning trials were 20 catch trials during which subjects moved to the point target but under veridical visual feedback. This experiment enabled us to ask if subjects are even able to adapt to two simultaneously imposed perturbations. This was important because we wished to ensure that any modulation in adaptation to the “rotation+gain” perturbation in Experiments 1 and 2 was not simply due to an inability to learn simultaneously imposed rotation and gain perturbations.

Note that in the main experiment, we did not include conditions where subjects adapted to task-relevant perturbations imposed alone (rotation perturbation while moving to the bar, and gain perturbation while moving to the arc). It was obvious that subjects would adapt to these perturbations; this has also been demonstrated by (Schaefer et al., 2012). Nonetheless, we still confirmed that this was indeed the case in a few pilot subjects (data not shown).

### Data Analysis

Hand position data was low pass filtered using a zero-phase lag Butterworth filter and differentiated to yield velocity values. Adaptation to the rotation was quantified by calculating the hand deviation from vertical normal on the catch trials, while that to the gain was quantified by assessing changes in initial movement extent. Hand deviation was calculated as the angle between the line connecting start position with the target and the line connecting start position with subject’s hand position at peak velocity. Initial movement extent was calculated as the distance between start position and hand position at peak velocity. We expected that a minifying gain would result in a greater initial displacement of the hand when compared to the “null” perturbation (Schaefer et al., 2012). One-way analysis of variance (ANOVA) was performed in Experiments 1 and 2 with condition (null, rotation, rotation+gain for Experiment 1; null, gain only, rotation+gain for Experiment 2) as the independent variable and hand deviation and initial displacement as the dependent variables (α=0.05). For Experiment 3, we assessed participants’ adaptation to the rotation+gain perturbation via changes in hand deviation and initial displacement on the catch trials relative to their own baseline trials using paired-samples t-tests. Post-hoc comparisons were done using Tukey-Kramer post-hoc tests. Trials on which subjects failed to initiate a movement or lifted the stylus off the tablet leading to loss of data, or made excessively slow (peak speed <5 cm/s) or fast (peak speed >100 cm/s) reaches were left out from our analyses. A total of 16 trials (8 catch trials from the first two experiments, and 3 baseline and 5 catch trials from experiment 3) were excluded across all our experiments based on the above criteria.

## RESULTS

### Experiment 1: Irrelevant Rotation Perturbation

In our first experiment, all reaches on the learning trials were made to the arc target alone. We compared catch trial performance across the ArcNull, ArcRot and ArcRot+Gain groups. Figure 2A shows the cursor trajectories of a representative subject from each of the groups on the catch trials. Unsurprisingly, for the ArcNull group, these trajectories remained relatively smooth and directed straight towards the target. In contrast, we observed a substantial clockwise deviation in the trajectory of the ArcRot group, which was a clear indication that subjects were adapting to the induced rotation. These deviations were corrected during the later portion of the movement to bring the cursor on to the target. Note that the rotation was irrelevant for the arc reaches, and in theory, even if left uncompensated, subjects would still be successful in reaching the arc target. However, it has been shown before that subjects cannot suppress adaptation to a rotation even when it is task-irrelevant (Schaefer et al., 2012). Our data for the ArcRot condition confirmed this trend. Remarkably though, for the ArcRot+Gain group, the angular deviation in trajectory was not observed, and subjects’ average trajectory was strikingly similar to the ArcNull subjects in terms of the direction (compare orange and black trajectories in Figure 2A, right). This suggested a notable reduction in adaptation to the rotation in the ArcRot+Gain group.

**Figure 2:**
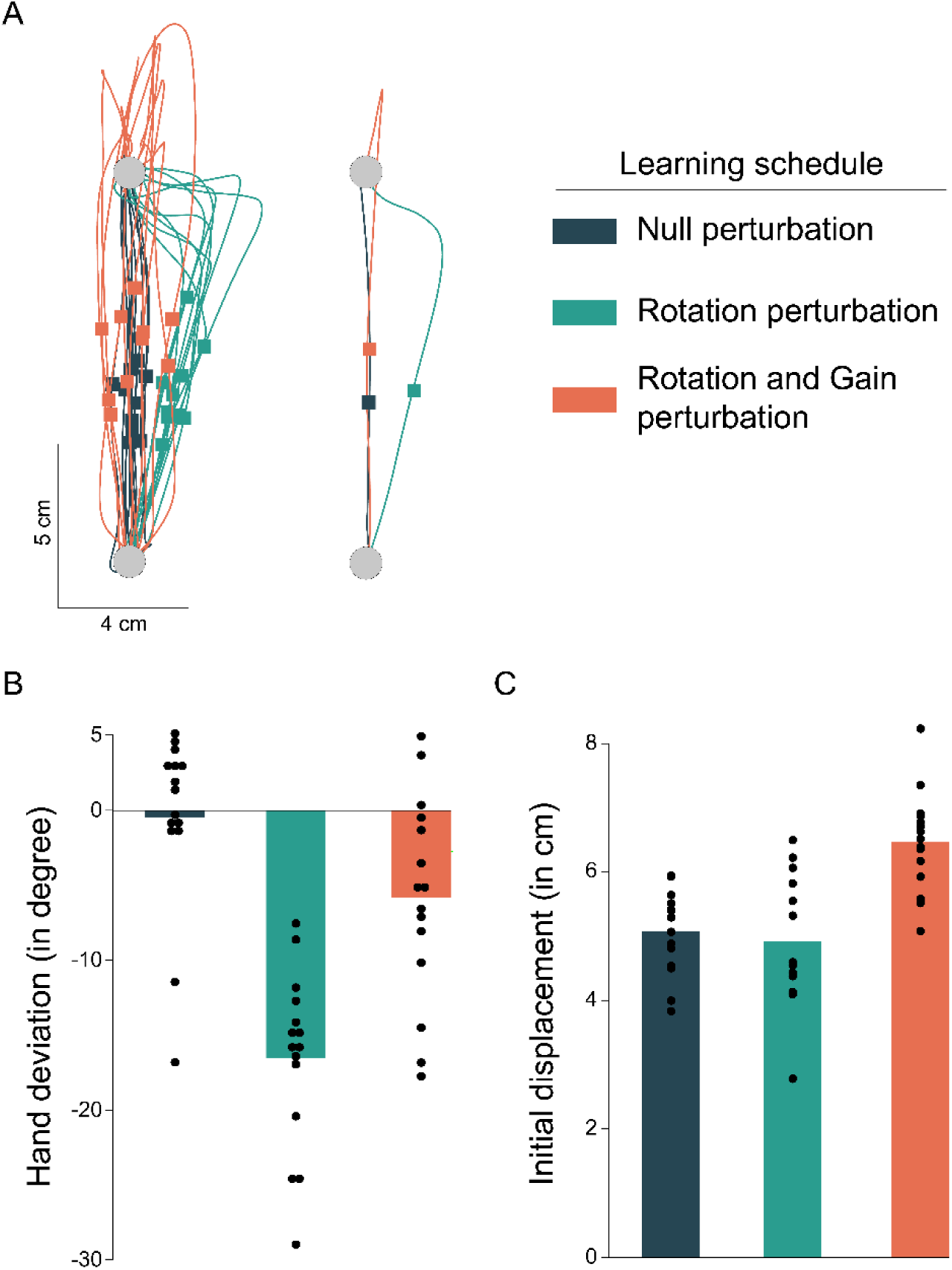
Adaptation while reaching to an arc target. (A) Hand trajectory data from a representative subject in each of the three perturbation conditions during the catch trials (right – all trials, left – averaged). Each square block indicates the hand position at peak velocity for the given trajectory. (B) Hand deviation at peak velocity for each of the three perturbation conditions (n=15 per condition). (C) Initial displacement at peak velocity for each condition during the arc target learning session. Each dot in (B) and (C) represents an individual subject’s data.

We then assessed how the ArcRot+Gain group adapted to the minifying gain. This was done by comparing the initial movement extent (i.e. displacement from movement initiation to peak velocity) of these subjects on the catch trials with that of the ArcNull group. In Figure 2A, the initial movement extent to peak velocity is shown using the colored squares on the respective trajectories. As is evident, the ArcRot+Gain subject covered a greater distance up to the time of peak velocity, indicating adaptation to the induced visuomotor gain. They also routinely overshot the target and then made corrective responses to bring the cursor back on to the target, again indicating that subjects were traveling farther to compensate for the reduced cursor gain.

Group level trends in hand deviation (Figure 2B) and movement extent (Figure 2C) were consistent with the patterns for individual subjects described above. Statistically, we observed a significant effect of group on the hand deviation of the catch trials (F(2,44) = 25.282, p < 0.001, ƞ^2^ = 0.55). Post hoc tests indicated a substantially higher deviation for the ArcRot group (mean ± SE: -16.544 ± 1.53⁰) compared to the ArcNull group (mean ± SE: -0.494 ± 1.55⁰, p<0.001). Notably though, hand deviation in the ArcRot+Gain group was drastically reduced (mean ± SE: -5.84 ± 1.77⁰, p<0.001, Figure 2B) falling nearly to the level observed for the ArcNull group (p = 0.063). There was also an effect of group on initial movement extent (Figure 2B; F(2,44) = 15.922, p<0.001, ƞ^2^= 0.43). Post hoc tests revealed that initial extent was not different between the ArcNull (mean ± SE: 5.082 ± 0.165 cm) and ArcRot (mean ± SE: 4.929 ± 0.26 cm) groups (p=0.868). However, the ArcRot+Gain subjects moved much farther (mean ± SE: 6.469 ± 0.2 cm) compared to both the ArcNull (p=0.001) and the ArcRot (p<0.001) groups.

To summarize, when reaching to the Arc target, subjects showed robust adaptation to a solo rotation perturbation even when it was irrelevant to the task goal. However, when paired with another relevant perturbation (gain), adaptation to the irrelevant rotation was selectively attenuated.

### Experiment 2: Irrelevant Gain Perturbation

In our second experiment, we probed whether a similar suppression would be evident when adapting to an irrelevant visuomotor gain paired with a relevant rotation. In this experiment, three groups of subjects (BarNull, BarGain, BarRot+Gain) reached to a bar target, which made the compensation of a gain perturbation irrelevant but a rotation perturbation highly relevant for task success. Figure 3A shows the hand trajectories of individual subjects in each of the three groups on the catch trials. Again, BarNull subjects showed canonical handpaths, directed smoothly and straight towards the target. Subjects in the BarGain group showed adaptation to the irrelevant gain perturbation when applied in isolation. We observed a clear increase in movement extent, represented as squares on the hand trajectories. That is, these subjects moved farther as they compensated for the gain. Notably however, subjects in the BarRot+Gain group did not show any scaling of movement extent; this is evident in the average trajectory (Figure 3A), in which the orange and black squares appear at nearly the same distance. Rather, these subjects seemed to only show a clear clockwise change in hand deviation in response to the simultaneously imposed (task-relevant) rotation.

**Figure 3:**
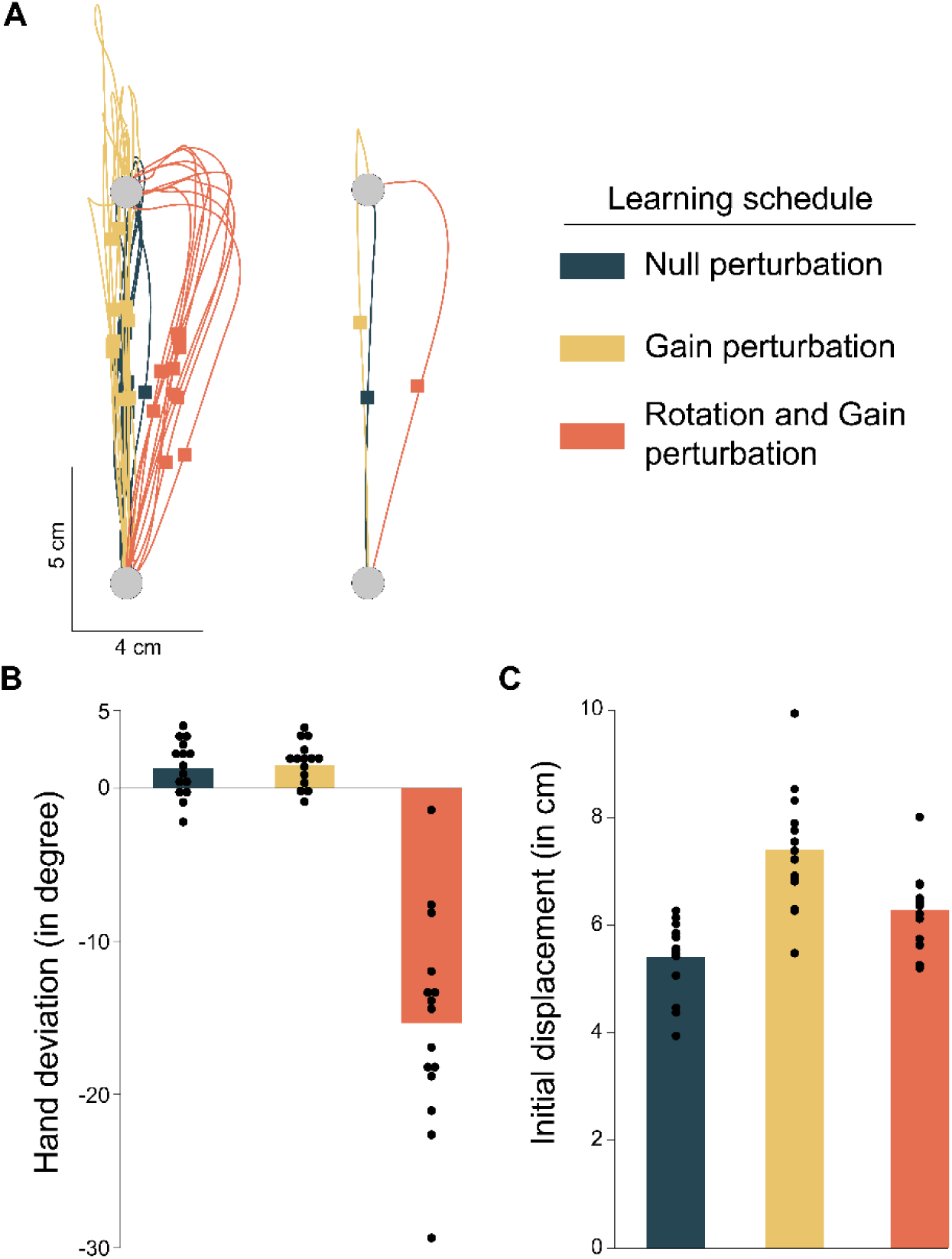
Adaptation while reaching to a bar target. (A) Hand trajectory data from a representative subject in each of the three perturbation conditions during the catch trials (right – all trials, left – averaged). Each square block indicates the hand position at peak velocity for the given trajectory. (B) Hand deviation at peak velocity for each of the three perturbation conditions (n=15 per condition). (C) Initial displacement at peak velocity for each condition during the bar target learning session. Each dot in (B) and (C) represents an individual subject’s data.

Group level data was very similar. We observed a clear main effect of group on the hand deviation. (Figure 3B; F(2,44)=82.66, p<0.001, ƞ^2^=0.794), with post hoc tests showing a robust increase for the BarRot+Gain group (mean ± SE: -15.37 ± 1.74⁰) compared to the BarGain (mean ± SE: 1.47 ± 0.37⁰, p<0.001) as well as the BarNull groups (mean ± SE: 1.27 ± 0.46⁰, p<0.001). Unsurprisingly, there were no significant differences in hand deviation between the BarGain and BarNull groups (p =0.99). A significant group effect was also found for movement extent (Figure 3B; F(2,44)= 21.44, p<0.001, ƞ^2^=0.505). Post hoc tests indicated that there was a robust increase in distance covered up to the time of peak velocity for the BarGain group (mean ± SE: 7.397 ± 0.278 cm) compared to the BarNull condition (mean ± SE: 5.407 ± 0.175 cm, p=0.002). However, when the gain perturbation was combined with the rotation (BarRot+Gain group), this movement extent was markedly diminished (mean ± SE: 6.277 ± 0.176 cm). Statistically, movement extent was significantly less in the BarRot+Gain subjects compared to the BarGain group (p<0.001), but the distance covered remained somewhat higher than the BarNull group (p=0.018). This suggested that while adaptation to the irrelevant gain perturbation was strongly attenuated, it perhaps wasn’t completely extinguished when a relevant rotation perturbation was simultaneously present. Note that no suppression was seen when the irrelevant gain perturbation was presented solo.

### Experiment 3: Control Experiment

Experiments 1 and 2 revealed a discernible reduction in adaptation to an irrelevant perturbation when it was paired with another relevant one. A pertinent question is whether this reduction was simply because the sensorimotor system is incapable of adapting to two simultaneously imposed rotation and gain perturbations. To assess this possibility, we conducted a third experiment in which subjects reached to a point target while experiencing the rotation+gain perturbation during learning. Note that in this case, both perturbations were task relevant. We then compared the performance on the catch trials of the learning block with that on baseline trials.

Figure 4A shows the hand trajectories of a representative subject from this group on baseline and catch trials. It is amply evident that this subject modified their movement direction to compensate for the rotation. Notably, this subject also showed adaptation to the gain; the initial movement extent was much greater on the catch trials compared to baseline as indicated by the difference in the location of the squares on the individual as well as average trajectories.

**Figure 4:**
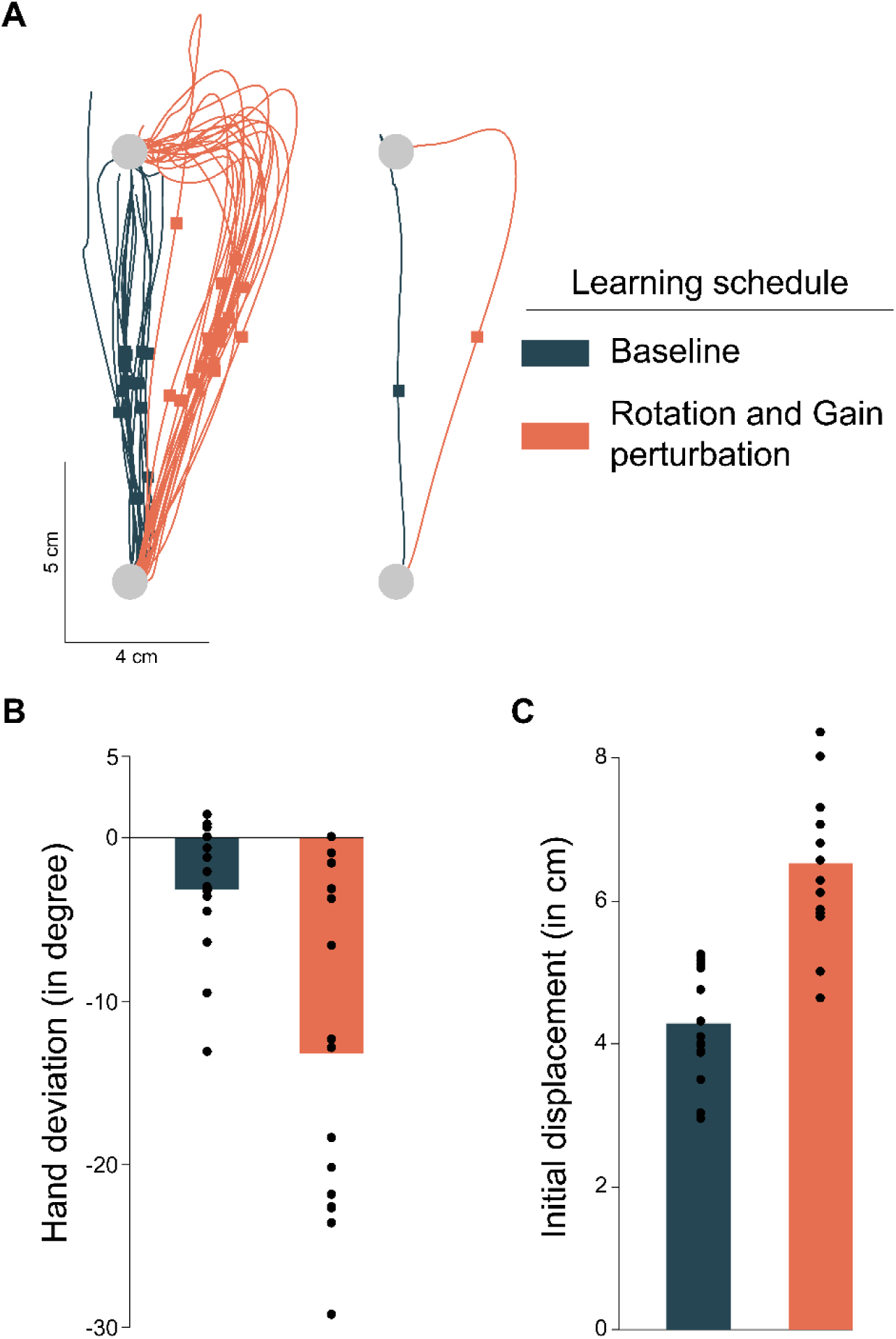
Adaptation while reaching a point target. (A) Hand trajectory data from a representative subject in baseline and the condition with two simultaneous perturbations during the catch trials (right – all trials, left – averaged). Each square block indicates the hand position at peak velocity for the given trajectory. While the actual rotation perturbation was induced in a clockwise direction and the adaptation observed in the counterclockwise direction, the adaptation is flipped here for homogeneous visualization purposes. (B) Hand deviation at peak velocity for the baseline and perturbation conditions (n=15). (C) Initial displacement at peak velocity for each condition during the point target learning session. Each dot in the (B) and (C) represents an individual subject’s data.

The same trend held at the group level. There was a robust increase in hand deviation (Figure 4B) as a consequence of adaptation to the rotation (mean ± SE: -13.353 ± 2.573°) when compared to same participants’ baseline trials (mean ± SE: -3.155 ± 1.034°; t(14)= -4.987, p<0.001). Likewise, there was an increase in movement extent (Figure 4C), reflecting adaptation to the visuomotor gain during the learning trials (mean ± SE: 6.525 ± 0.264 cm) relative to baseline (mean ± SE: 4.284 ± 0.202 cm; t(14)= 6.276, p<0.001). These results were a clear indication that the sensorimotor system is indeed capable of adapting to two simultaneously imposed perturbations if they are relevant for task success.

## DISCUSSION

Previous studies have proposed that motor adaptation in the presence of prediction errors is obligatory and cannot be suppressed (Mazzoni & Krakauer, 2006; Morehead et al., 2017). Contrary to this, we observed specific and robust attenuation of adaptation in response to errors that were irrelevant to achievement of the task goal. This attenuation, however, occurred only when multiple feedback signals were present simultaneously, i.e. when there was a task-relevant error present with an irrelevant error. Indeed, like past work, adaptation to relevant or irrelevant errors occurring alone could not be attenuated. These results further nuance our understanding of error-driven motor adaptation and provide support to the idea that adaptation can indeed be modulated in a task-dependent manner.

Past work has suggested that the motor system responds differently to errors of different sizes likely because it assigns these errors to different sources (Kim et al., 2019; Morehead et al., 2017), and that this response likely saturates for large errors (Hutter & Taylor, 2018; Morehead et al., 2017). An error of a fixed magnitude has also been shown to have some influence on the ensuing motor behavior depending on its relevance to the task goal. This, in fact, appears to be the case for both between-trial adaptation and within-trial feedback control. Schaefer et al. (2012), for instance, examined the strength of the after-effects on catch trials when subjects underwent adaptation in the presence of sensory prediction errors that were either task relevant or irrelevant. They found smaller after-effects in the task-irrelevant condition, suggesting that the adaptive response was modulated by error relevance. Likewise, in an elegant study, Reichenbach et al. (Reichenbach et al., 2014) compared the magnitude of corrective responses when subjects made point to point reaching movements under two conditions: first, when task relevant errors were introduced by displacing either a cursor that represented the hand position or displacing the target of the reach, and second, when task irrelevant errors were introduced by displacing distractors around the target or the cursor. They found that corrective responses were much stronger when the errors were task relevant versus when they were not; similar relevance-sensitivity was seen in a force-field paradigm (Heald et al., 2018). What is most interesting in both these sets of results, however, is the finding that neither the corrective nor the adaptive responses were suppressed when the errors were irrelevant. In line with these results, Morehead et al. (2017) demonstrated using error feedback clamped to a certain direction independent of hand movement and clear instructions to ignore this task irrelevant error, that subjects showed clear after-effects similar to those seen following implicit adaptation to task relevant errors. This is where our current findings depart from these previous results; we show a near complete absence of adaptation under task irrelevant conditions. Importantly however, this suppression was observed only when multiple error signals were present simultaneously. In the presence of a single error, our findings align with past work in that we observe adaptation even when the error is irrelevant.

In this study, the task relevant and irrelevant errors were induced using either rotation or gain perturbations, which bring about errors in direction and extent respectively. A large body of prior work has argued that these two movement parameters are independently specified in the brain during the movement planning stages (Favilla & De Cecco, 1996; Krakauer et al., 1995, 2004; Larish & Frekany, 1985; Messier & Kalaska, 1997). In particular, it is thought that movement direction is planned in a feedforward manner while the control of movement extent requires sensory feedback (Larish & Frekany, 1985; Riehle & Requin, 1989). Correspondingly, cells in premotor and motor cortex, the activity of which correlates with initial movement direction, final position and movement amplitude have been identified (Riehle & Requin, 1989). In these cells, firing patterns correlate well with movement direction before movement onset while correlations with movement amplitude evolve during the movement, suggesting independent control of these two aspects of movement. Our results are in agreement with this distinction and show that just like direction and extent are separately specified, *errors* in direction and extent are also independently processed in the brain. This potentially enables the brain to independently establish scaling factors that relate movement amplitude to peak velocity as required for gain adaptation, and reference axes as required for direction adaptation. This is in line with prior work that established differential generalization patterns following rotation or gain adaptation, thereby arguing for differential representation of these parameters in neural circuits (Krakauer et al., 1995; Messier & Kalaska, 1997). One could thus argue that this neural separation is the likely basis of the selective attenuation of adaptation to rotation and gain perturbations.

However, recent results suggest that attenuation is likely to occur even when the feedback signals are related to just one of these parameters alone. Tsay et al. (2024) very recently demonstrated that when subjects were exposed to two simultaneous cursor rotations, one relevant and another irrelevant to the task goal, there was an overall reduction in the magnitude of adaptation compared to a single rotated cursor condition. Notably, the position of the irrelevant cursor was inconsequential, pointing to a largely non-specific effect that occurred likely due to the presence of a distracting stimulus (the irrelevant cursor). A stronger suppression was observed by Kasuga et al. (2013), who showed neither an after-effect nor any generalization when subjects “adapted” to an irrelevant rotation presented simultaneously with a relevant one, suggesting that learning could be entirely suppressed. These results indicate that selective suppression may not be related to the type of perturbation (rotation versus gain), but perhaps has a different basis.

One candidate explanation for this may then be that the nervous system selectively filters out task irrelevant information in a manner akin to that in the biased competition model of visual attention (Desimone & Duncan, 1995). In the biased competition framework, the most salient or task-relevant attributes of a visual stimulus are prioritized when task-irrelevant information is simultaneously present and competes for neural resources with limited processing capacity. This then acts as an input to the attentional system so that the biased stimulus can be attended to. Analogously, for adaptation, relevance of the error to the task goal may be the salient attribute that affords it priority over the irrelevant error for subsequent processing and adaptive changes. Importantly, this does not imply that the irrelevant error is not monitored; indeed, in the results of Tsay et al. (2024), it was shown that subjects who attended to an irrelevant target/cursor pair also demonstrated small after-effects indicating some adaptation despite a task relevant error being present. Nonetheless, their work and our current results suggest that given limited processing capacity, the task relevant error is prioritized. When the irrelevant error occurs alone, the competition with the relevant attribute does not exist, and this error can alone bring about adaptive changes.

In sum, our study provides robust evidence to support the notion that adaptation to sensory prediction error can be strongly attenuated depending upon the relevance of the error. However, this attenuation of adaptation seems to occur only when multiple error signals are present simultaneously. In fact, when presented in isolation, the presence of the error signal is sufficient to elicit adaptation regardless of its relevance to the task success. These findings further nuance the classical view that adaptation to sensory prediction error is obligatory in nature.

## ACKNOWLEDGEMENTS

This work was supported by funding from the Department of Science and Technology, Government of India to PKM. Support from IIT Gandhinagar is also acknowledged. The authors thank Sankha Subhra Bhattacharjee for assistance with data collection, and Adarsh Kumar and Gaurav Panthi for assistance with data analysis and discussions on this topic.

## CONFLICT OF INTEREST

None

## Notes

### Competing Interest Statement

The authors have declared no competing interest.

